# *NANOG* is dispensable for hypoblast but essential for embryonic disc development during conceptus elongation in sheep

**DOI:** 10.64898/2026.09.24.754098

**Authors:** Nuria Martínez de los Reyes, Melissa Carvajal-Serna, Pilar Marigorta, Adolfo Toledano-Díaz, Julián Santiago-Moreno, Pablo Bermejo-Álvarez, Priscila Ramos-Ibeas

**Author notes:** **Correspondence**: Animal Reproduction Department, Instituto Nacional de Investigación Agraria y Alimentaria, Consejo Superior de Investigaciones Científicas, Avenida Puerta de Hierro 18, 28040, Madrid, Spain.

## Abstract

*NANOG* is a conserved regulator of epiblast identity and is essential for hypoblast development and embryo survival after the blastocyst stage in mice. However, its role during post-blastocyst development remains poorly understood in non-rodent mammals. Here, we investigated the function of *NANOG* during ovine embryogenesis using CRISPR/Cas9-mediated gene ablation. *NANOG* ablation did not affect blastocyst formation or the initial development of the epiblast. However, *NANOG*-deficient blastocysts contained significantly fewer SOX17+ hypoblast cells by day (D) 8, followed by reduced hypoblast migration during early post-hatching stages in D12 *in vitro* embryos. These defects were transient, as hypoblast development was comparable between *NANOG*-deficient and wild-type conceptuses during subsequent *in vivo* conceptus elongation from embryonic day (E) 11 onwards. In contrast, although extraembryonic membranes development and conceptus elongation proceeded normally, NANOG-deficient conceptuses lacked SOX2+ epiblast cells and failed to form an embryonic disc (ED) from E11 onwards. These findings demonstrate that *NANOG* is dispensable for hypoblast development but is required for epiblast maintenance and ED formation during conceptus elongation in sheep.

## 2. Introduction

During early mammalian embryo development, two critical and consecutive lineage segregation events occur. The first results in the formation of the inner cell mass (ICM) and trophectoderm (TE) during blastocyst development (Niwa *et al*., 2005). Subsequently, the second lineage segregation event differentiates the ICM into the epiblast and hypoblast, also referred to as the primitive endoderm (PrE) (Artus *et al*., 2011, Plusa *et al*., 2008). The epiblast gives rise to the embryo proper, whereas the hypoblast and TE contribute to the formation of the extra-embryonic membranes (EEMs), including the yolk sac and placenta, respectively (Martínez de los Reyes *et al*., 2026).

The molecular mechanisms regulating these lineage specification events, and the transcription factors involved, have been extensively studied in the mouse model. However, an increasing number of studies have revealed substantial interspecies differences in early mammalian embryonic development. In mice, the first lineage segregation is regulated by the Hippo signalling pathway, which is influenced by cell polarity and position at the morula stage. In outer cells, Hippo signalling is inactive, resulting in the nuclear accumulation of Yes-associated protein (YAP). Nuclear YAP interacts with TEAD4 to activate the expression of TE-specific genes, including *Cdx2* and *Gata3*. In contrast, Hippo signalling is active in inner cells, leading to YAP phosphorylation and cytoplasmic retention. Under these conditions, TEAD4 is not expressed, and ICM-specific genes are upregulated, including *Oct4* and *Sox2* (Nishioka *et al*., 2009, Nishioka *et al*., 2008, Ralston *et al*., 2010). Notably, the role of TEAD4 as a master regulator of TE differentiation and blastocyst formation in mice is not conserved across all mammals, including rabbits, cattle, and humans (Pérez-Gómez *et al*., 2024, Pérez-Gómez *et al*., 2021, Stamatiadis *et al*., 2022, Wu *et al*., 2024). Similarly, although ablation of *Cdx2*, a downstream target of TEAD4, prevents blastocyst formation in mice (Strumpf *et al*., 2005, Wicklow *et al*., 2014), bovine embryos lacking *CDX2* can form blastocysts (Shi *et al*., 2023). Also, sheep and cattle embryos lacking *GATA3* are able to form blastocysts (Martínez de Los Reyes *et al*., 2025, Shi *et al*., 2023).

The second lineage segregation event in mice is initially driven by the mutually exclusive expression of the transcription factors NANOG and GATA6 within ICM cells, which are arranged in a “salt-and-pepper” pattern. These cells subsequently segregate to form the epiblast and PrE or hypoblast, respectively (Artus *et al*., 2011, Plusa *et al*., 2008). In addition to its role in epiblast vs. hypoblast specification, NANOG is essential for maintaining pluripotency (Mitsui *et al*., 2003). Epiblast-specific NANOG expression is conserved across multiple mammalian species, including mice, ungulates (González-Brusi *et al*., 2026, Kuijk *et al*., 2012, Martínez de Los Reyes *et al*., 2024, Ramos-Ibeas *et al*., 2019), and primates (Nakamura *et al*., 2016, Petropoulos *et al*., 2016), although its timing differs among species. In mouse embryos, NANOG expression is initiated at the morula stage and subsequently becomes restricted to the epiblast upon blastocyst formation (Bessonnard *et al*., 2014, Plusa *et al*., 2008). In contrast, in human, bovine and ovine embryos, NANOG expression is first detected in the ICM of early blastocysts (Blakeley *et al*., 2015, Cauffman *et al*., 2009, Kuijk *et al*., 2012, Madeja *et al*., 2013, Martínez de Los Reyes *et al*., 2024, Petropoulos *et al*., 2016). In mice, *Nanog* ablation disrupts epiblast specification and results in developmental arrest before embryonic day (E) 5.5, shortly after implantation (Frankenberg *et al*., 2011, Messerschmidt and Kemler, 2010, Mitsui *et al*., 2003). Although *Nanog*-null blastocysts initially express GATA6 in the ICM, expression of later hypoblast markers, including SOX17 and GATA4, is impaired (Bower *et al*., 2026, Frankenberg *et al*., 2011, Messerschmidt and Kemler, 2010). However, in human embryos, *NANOG* ablation did not affect hypoblast development, highlighting differences from the mouse (Bower *et al*., 2026). In pig embryos, *NANOG* ablation results in reduced numbers of OCT4- and SOX2-positive cells, but no evident differences in the number of hypoblast cells were observed (Lee *et al*., 2023). In bovine embryos, contrasting results have been reported: one study showed reduced *GATA6* and *SOX2* mRNA expression following *NANOG* ablation (Ortega *et al*., 2020), whereas another reported normal numbers of SOX2- and OCT4-positive ICM cells and no apparent effect on hypoblast formation (Springer *et al*., 2021). Collectively, these findings suggest species-specific differences in the role of NANOG in hypoblast development, and its function in ungulate embryos remains to be fully elucidated. Unfortunately, studies in non-rodent species have largely been limited to the blastocyst stage, impeding to evaluate further hypoblast and epiblast development.

Whereas mouse and primate embryos implant shortly after blastocyst hatching from the *zona pellucida*, ungulate embryos undergo an extended preimplantation period known as conceptus elongation. Following hatching, the hypoblast proliferates and migrates along the inner surface of the TE, and both EEMs will undergo extensive proliferation. Concurrently, epiblast cells proliferate and organize into a compact embryonic disc (ED), which subsequently undergoes symmetry breaking and gastrulation before implantation (Brooks *et al*., 2014, Maddox-Hyttel *et al*., 2003, Perez-Gomez *et al*., 2021). The role of NANOG during this developmental window in ungulates remains unknown. We previously developed an extended culture system that supports post-hatching development, including complete hypoblast migration along the TE and ED development, in both bovine and ovine embryos (Ramos-Ibeas *et al*., 2022, Ramos-Ibeas *et al*., 2020, Ramos-Ibeas *et al*., 2023). This system provides an *in vitro* platform for functional studies of molecular mechanisms regulating post-hatching development.

The aim of this study was to elucidate the role of *NANOG* in ovine embryonic development using CRISPR/Cas9-mediated gene ablation and complementary *in vitro* and *in vivo* approaches. Specifically, we investigated the effects of NANOG ablation on epiblast and hypoblast differentiation at the blastocyst stage, as well as during post-hatching stages *in vitro* and conceptus elongation *in vivo*.

## 3. Materials and methods

### 3.1. Ethics statement

All experimental procedures were approved by the INIA Animal Care Committee and the Madrid Regional Authorities (PROEX 059.2/22) in accordance with European legislation.

### 3.2. In vitro production of NANOG knock-out (KO) ovine embryos

Ovine *NANOG* gene was disrupted by introducing frame-disrupting insertions or deletions (indels) generated during non-homologous end joining (NHEJ) repair of CRISPR/Cas9-induced double-strand breaks (DSB). The online software CRISPOR (Concordet and Haeussler, 2018) was used to design a single guide RNA (sgRNA) targeting exon 2 of the *NANOG* gene. The sgRNA targeting the sequence CAGAGTGAAACCACTGTCCC was produced using the Guide-it sgRNA In Vitro Transcription Kit (Takara) with a primer containing the T7 promoter and the sgRNA sequence (**Table 1**). Capped polyadenylated Cas9 messenger RNA (mRNA) was transcribed *in vitro* from the plasmid pMJ920 (Addgene 42234) using the mMESSAGE mMACHINE T7 ULTRA kit (Life Technologies) and purified using the MEGAClear kit (Life Technologies).

**Table 1.**
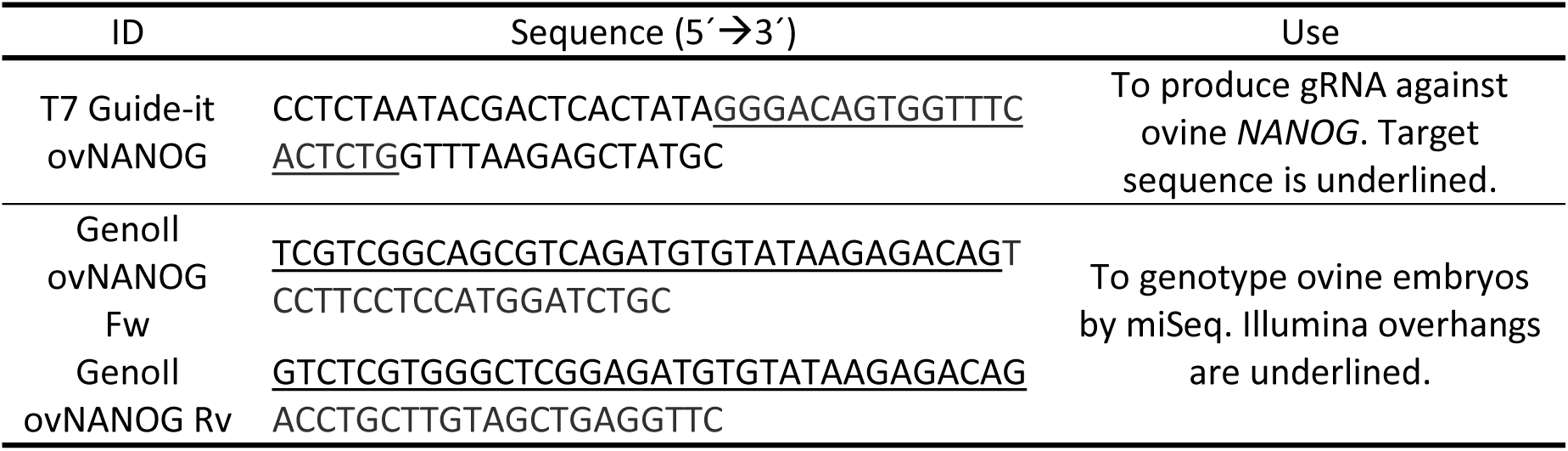
Details of primers used for guide RNA (gRNA) production, as well as for genotyping.

Cumulus-oocyte complexes (COCs) were aspirated from 2 to 8 mm diameter follicles of ovine ovaries collected at a local slaughterhouse using a 21 G needle connected to an aspiration pump (VMAR 5100, Cook) set to −25 mmHg. Aspirates were collected in a 50 ml tube containing Euroflush medium (IMV Technologies). COCs exhibiting compact cumulus and homogeneous cytoplasm were selected and matured for 22-24 h in TCM-199 supplemented with 40 μg/ml gentamicin sulphate, 10 % (v/v) fetal bovine serum (FBS), and 10 ng/ml epidermal growth factor (EGF) at 38.5 °C in an atmosphere of 5 % CO_2_ with maximum humidity. Following 22 h maturation, cumulus cells were removed by vortexing for 3 min in 1 ml phosphate-buffered saline (PBS) containing 300 µg/ml hyaluronidase. Cytoplasmic microinjection was then performed using a spike-end micropipette (5 µm internal diameter; BioMedical Instruments, Germany) connected to a manual hydraulic microinjector (CellTram Air, Eppendorf, Germany) under a Nikon Eclipse TE300 microscope.

Oocytes were randomly assigned to two groups: 1) *NANOG*-targeted group, microinjected with 300 ng/µl Cas9-encoding mRNA and 100 ng/µl sgRNA against *NANOG* (∼2/3 of the oocytes per replicate), and 2) control group, microinjected with 300 ng/µl Cas9-encoding mRNA alone (∼1/3 of the oocytes per replicate). The control group served as a microinjection control where embryos are unedited.

Immediately after microinjection, oocytes were fertilized in groups of approximately 25 using frozen-thawed ram sperm purified with Bovi-Pure® (Nidacon) at a final concentration of 2 × 10^6^ spermatozoa/ml. Gametes were co-incubated in 50 µl droplets of IVF medium (Stroebech media, Denmark) supplemented with 10 % (v/v) heat-inactivated oestrous sheep serum and 50 IU/ml heparin, covered with mineral oil, at 38.5 °C in an atmosphere of 5 % CO_2_ with maximum humidity. Semen from the same ram was used for all experimental replicates to avoid potential confounding effects of paternal variability on embryo development.

At 18 h post-insemination, presumptive zygotes were denuded by manual pipetting and cultured in groups of approximately 25 in 50 μl droplets of IVC medium (Stroebech media, Denmark), covered with mineral oil, at 38.5 °C in a humidified atmosphere of 5 % CO_2_, 5 % O_2_, and 90 % N_2_. Cleavage was assessed at 48 h, and blastocyst rates were recorded at day (D) 8.

### 3.3. Post-hatching development system

On D6 and D7 post-fertilization, blastocysts were transferred to SPL3D^TM^ Cell Floater low-attachment dishes (SPL Life Sciences, Korea) and cultured in N2B27 medium [1:1 Neurobasal and DMEM/F12 medium supplemented with penicillin/streptomycin, 2 mM L-glutamine, and N2 and B27 supplements (Thermo Fisher Scientific)] supplemented with 20 ng/ml Activin A and 10 µM ROCK inhibitor Y-27632 (StemCell Technologies), as previously described (Ramos-Ibeas *et al*., 2022) at 38.5 **°**C in a water-saturated atmosphere of 5 % CO_2_, 5 % O_2_, and 90 % N_2_. Half of the culture medium was replaced every other day until D12, when pictures were taken and embryos were fixed.

### 3.4. Development of NANOG KO conceptuses in vivo

D7 blastocysts from the *NANOG*-targeted group (partially composed of KO embryos) and the control group (composed of wild-type [WT] embryos only) were transferred to four synchronized recipient ewes. Estrous synchronization was achieved by inserting a 0.35 g progesterone-releasing device (CIDR-Ovis, Zoetis). Cloprostenol (100 µg, Estrumate®, MSD) was administered 11 days after CIDR insertion, and PMSG (450 IU, Sincropart PMSG, CEVA) was injected on the day of CIDR removal (12 days post-insertion). Embryo transfer was performed 8 days after CIDR removal. The presence of corpora lutea was confirmed by abdominal laparoscopy, and embryos were transferred into the cranial section of the uterine horns. After 5, 6, or 7 days of development *in vivo*, ewes were slaughtered and embryos were recovered by uterine flushing with warmed recovery medium (Euroflush®, IVM Technologies). Conceptus length was measured, and embryos were fixed for further analyses.

### 3.5. Immunofluorescence analysis of lineages development

If present, the *zona pellucida* was removed from unhatched D8 blastocysts by brief incubation in PBS (pH 2) at 37 **°**C. Embryos were fixed in 4 % paraformaldehyde (PFA) for 15 min at RT and washed in PBS containing 1 % bovine serum albumin (BSA). When immunofluorescence was not performed immediately after fixation, embryos were stored at 4 **°**C in PBS-1 % BSA and incubated in 125 mM glycine in PBS-1 % BSA for 3 hours at 4 **°**C prior to immunostaining. Permeabilization was performed using 1 % Triton X-100 in PBS for 40 min at room temperature (RT), followed by blocking in PBS containing 10 % FBS and 0.02 % Tween-20 for 1 hour at RT. Embryos were then incubated overnight at 4 **°**C with primary antibodies to detect epiblast and hypoblast (**Table 2**). After four washes of 10 min in PBS-1 % BSA, embryos were incubated with the corresponding secondary antibodies (**Table 2**), and counterstained with DAPI for 1 hour at RT. Finally, embryos were washed four times in PBS-1% BSA. To acquire three-dimensional images, embryos were placed in PBS-1% BSA microdrops delimited with a PAP pen (Kisker Biotech GmbH) on a coverslip. Microdrops were covered with an Invitrogen™ CoverWell™ Incubation Chamber Gasket (Life Technologies) to prevent embryo compression and to allow embryo recovery for genotyping after imaging. Embryo images were acquired every 5 µm along the Z-axis using a structured illumination system consisting of a Zeiss Axio Observer microscope coupled to ApoTome.2.

**Table 2.**
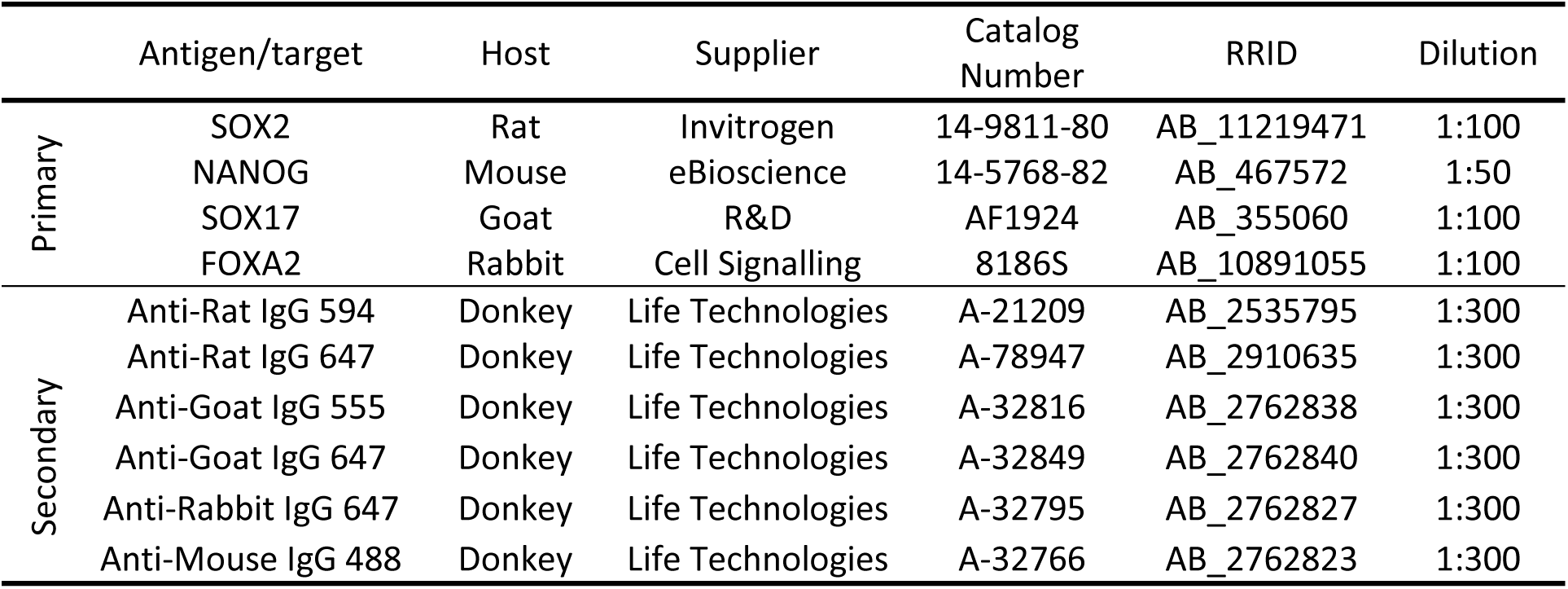
Details of the primary and secondary antibodies used. Antigens targeted by the primary antibodies and descriptions of the secondary antibodies, as well as host species, catalogue numbers, RRID (Research Resource Identifier), and dilutions used for each antibody are provided.

|  | Antigen/target | Host | Supplier | Catalog Number | RRID | Dilution |
| --- | --- | --- | --- | --- | --- | --- |
| Primary | SOX2 | Rat | Invitrogen | 14-9811-80 | AB_11219471 | 1:100 |
|  | NANOG | Mouse | eBioscience | 14-5768-82 | AB_467572 | 1:50 |
|  | SOX17 | Goat | R&D | AF1924 | AB_355060 | 1:100 |
|  | FOXA2 | Rabbit | Cell Signalling | 8186S | AB_10891055 | 1:100 |
| Secondary | Anti-Rat IgG 594 | Donkey | Life Technologies | A-21209 | AB_2535795 | 1:300 |
|  | Anti-Rat IgG 647 | Donkey | Life Technologies | A-78947 | AB_2910635 | 1:300 |
|  | Anti-Goat IgG 555 | Donkey | Life Technologies | A-32816 | AB_2762838 | 1:300 |
|  | Anti-Goat IgG 647 | Donkey | Life Technologies | A-32849 | AB_2762840 | 1:300 |
|  | Anti-Rabbit IgG 647 | Donkey | Life Technologies | A-32795 | AB_2762827 | 1:300 |
|  | Anti-Mouse IgG 488 | Donkey | Life Technologies | A-32766 | AB_2762823 | 1:300 |

### 3.6. Embryo genotyping by deep sequencing

CRISPR/Cas9-induced indels were identified by deep sequencing (MiSeq, Illumina, San Diego, USA) as previously described (Lamas-Toranzo *et al*., 2019). After fluorescence image acquisition, D8 and D12 *in vitro* embryos and fragments of elongated conceptuses were placed at the bottom of 0.2 ml PCR tubes and stored at −20 °C. D8 blastocysts were lysed in 8 µl and D12 *in vitro* embryos or conceptus fragments in 10 µl of Arcturus® Picopure® extraction buffer (Thermo Fisher Scientific) at 65°C for 1 hour, followed by inactivation at 95°C for 10 min. A genomic sequence containing the sgRNA target sequence was amplified by PCR using primers containing Illumina adaptors (**Table 1**) and GoTaq® DNA polymerase (Promega, Madison, USA). PCR conditions were: 96°C for 2 min; 30 cycles of 96°C for 20 seconds, 64°C for 30 seconds, and 72°C for 30 seconds; followed by a final extension at 72°C for 5 minutes. PCR products were purified using AMPure XP Beads (Beckman Coulter, USA) at a beads-to-sample ratio of 0.8:1. Indexed libraries were generated in a second PCR using Nextera XT Index Kit v2 primers (Illumina, San Diego, USA) with the following conditions: 95°C for 3 minutes; 8 cycles of 95°C for 30 seconds, 55°C for 30 seconds, 72°C for 30 seconds; and a final extension at 72°C for 5 minutes. PCR products were purified with AMPure XP beads at a beads-to-sample ratio of 1.12:1. Finally, libraries from different embryos were pooled at 8 nM and sequenced on an Illumina MiSeq system by an external company. Sequencing reads were aligned to the WT reference sequence using the *BWA-MEM* aligner (v.0.7.17, (Li and Durbin, 2010)), then sorted and indexed using the *SAM tools* package (v.1.16.1 (Li *et al*., 2009)). Variant calling was performed with FreeBayes (Garrison and Marth, 2012). VCF files were processed with an in-house script to select only potential indels within the CRISPR/Cas9 target region that passed the quality control. Results were visually confirmed using the Integrative Genomics Viewer (Thorvaldsdóttir *et al*., 2013).

Based on the analysis of ˃1,000 reads of the target region per embryo, embryos were classified as: WT; containing no mutated alleles or KO; containing only frame-disrupting indels (indels not divisible by three). Edited embryos containing at least one non-frame-disrupting indel were excluded from the analyses.

### 3.7. Experimental design and end-point analyses

In a first experiment (5 experimental replicates), we assessed the role of NANOG during blastocyst development. Blastocyst rates from *NANOG*-targeted and control groups were recorded at D8, and lineages development was analysed by immunofluorescence using ICM/epiblast (SOX2 and NANOG) and hypoblast (SOX17) markers. Total cell numbers and the number of cells expressing each marker were quantified using the multi-point counter plugin in ZEN 3.2 software (Carl Zeiss Microscopy GmbH, Germany). Embryos from *NANOG*-targeted group were subsequently genotyped.

In a second experiment (3 experimental replicates), we characterized the role of NANOG during post-hatching development *in vitro*. Embryos were cultured until D12, photographed using a stereomicroscope (Zeiss Stemi 305), and embryo area was measured using ZEN 3.2 software (Carl Zeiss Microscopy GmbH, Germany). Embryo survival was assessed based on morphological criteria: alive embryos were able to maintain the blastocoel, whereas dead embryos collapsed. Surviving embryos were analysed for lineages development by immunofluorescence for epiblast (SOX2 and NANOG) and hypoblast (SOX17) markers. Epiblast survival was defined by the presence of SOX2+ cells at the end of the culture period and embryonic disc (ED) formation as the presence of a compact group of ≥ 30 SOX2+ cells. SOX2+ and NANOG+ cell number were manually counted using the multi-point counter plugin in ZEN 3.2 (Carl Zeiss Microscopy GmbH, Germany). Hypoblast migration was quantified as the extent of SOX17+ hypoblast cell coverage along the inner surface of the TE. Each spherical embryo was bisected along the Z-axis, and orthogonal projections were produced. Total area and the area covered by SOX17+ hypoblast cells were determined for each projection, and hypoblast migration was calculated as the average proportion of hypoblast-covered area relative to the total area from both projections, divided by the total area. Then, embryos from *NANOG*-targeted group were genotyped.

In a third experiment (4 experimental replicates), the role of NANOG during conceptus elongation *in vivo* was investigated. Conceptuses recovered at embryonic days (E) 11, 12, and 13 were fixed, imaged and measured using Fiji software (Schindelin *et al*., 2012), and immunostained for epiblast (SOX2 and NANOG) and hypoblast (SOX17 and FOXA2) markers. Hypoblast migration was considered complete when the entire inner surface of the conceptus was covered by SOX17+ or FOXA2+ cells. Embryonic disc formation was assessed morphologically and confirmed by SOX2 staining. Embryos were then genotyped.

The number of microinjected oocytes and embryos developed per experiment is shown in **Table 3**.

**Table 3.**
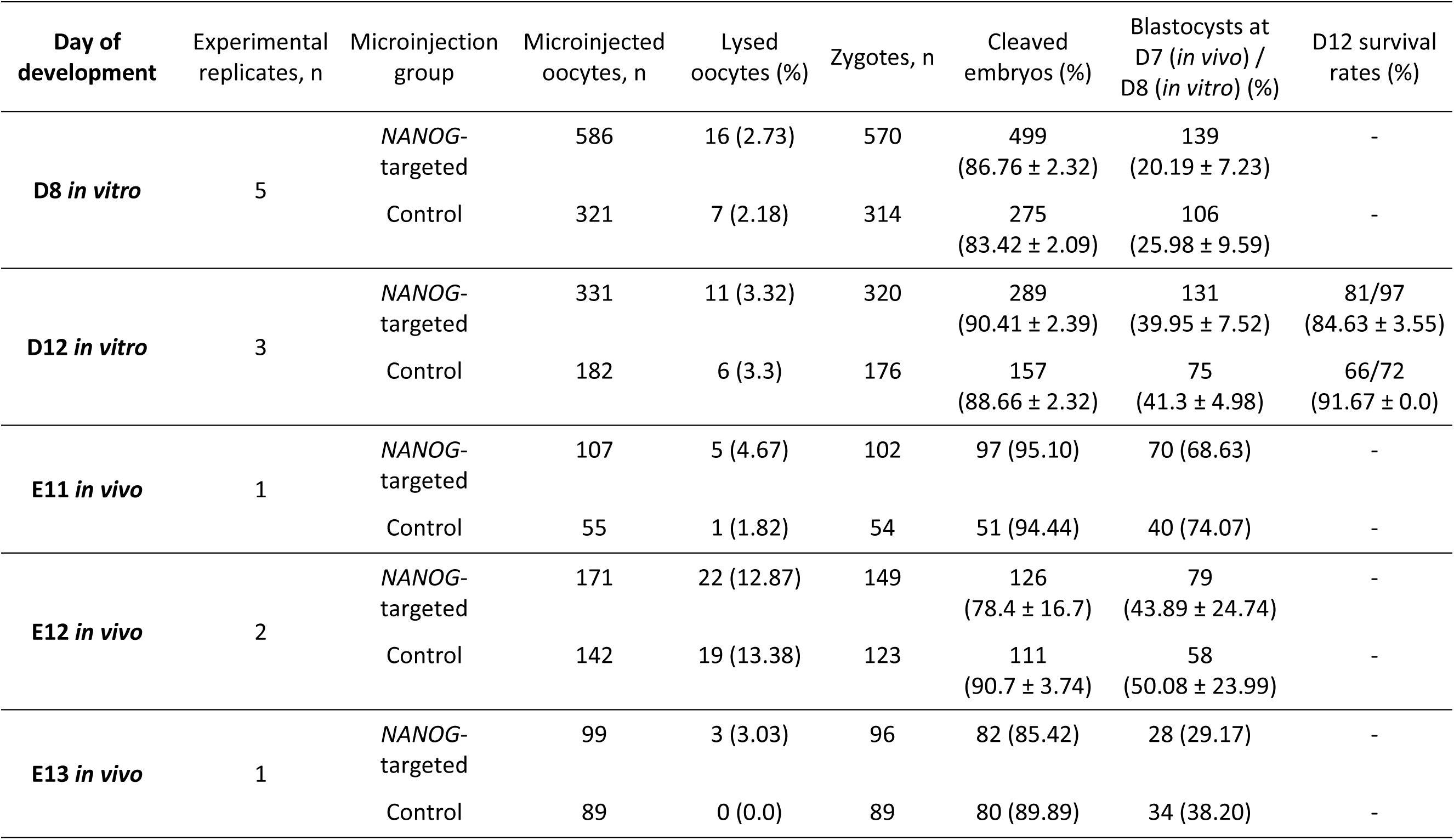
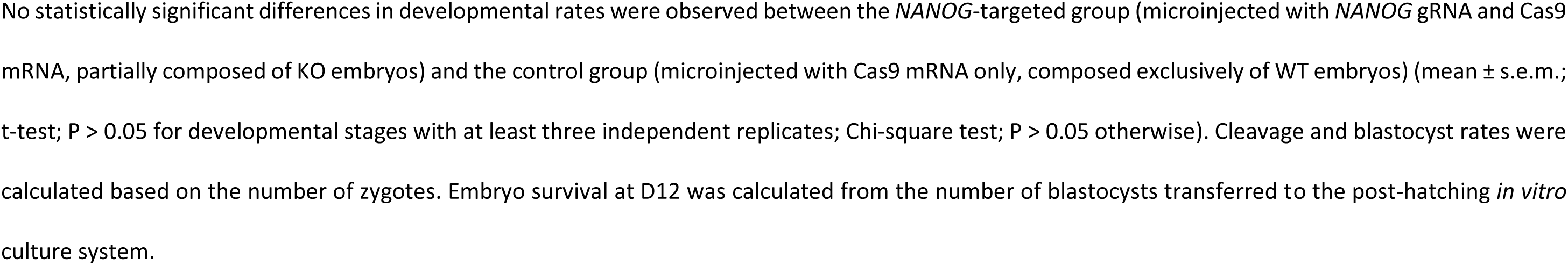
Number of embryos and developmental rates in each experimental group at day (D) 8 and 12 of *in vitro* development, and at embryonic days (E) 11, 12, and 13 of *in vivo* development.

| Day of development | Experimental replicates, n | Microinjection group | Microinjected oocytes, n | Lysed oocytes (%) | Zygotes, n | Cleaved embryos (%) | Blastocysts at D7 (*in vivo*) / D8 (*in vitro*) (%) | D12 survival rates (%) |
| --- | --- | --- | --- | --- | --- | --- | --- | --- |
| **D8 *in vitro*** | 5 | *NANOG*-targeted | 586 | 16 (2.73) | 570 | 499<br>(86.76 ± 2.32) | 139<br>(20.19 ± 7.23) | - |
|  |  | Control | 321 | 7 (2.18) | 314 | 275<br>(83.42 ± 2.09) | 106<br>(25.98 ± 9.59) | - |
| **D12 *in vitro*** | 3 | *NANOG*-targeted | 331 | 11 (3.32) | 320 | 289<br>(90.41 ± 2.39) | 131<br>(39.95 ± 7.52) | 81/97<br>(84.63 ± 3.55) |
|  |  | Control | 182 | 6 (3.3) | 176 | 157<br>(88.66 ± 2.32) | 75<br>(41.3 ± 4.98) | 66/72<br>(91.67 ± 0.0) |
| **E11 *in vivo*** | 1 | *NANOG*-targeted | 107 | 5 (4.67) | 102 | 97 (95.10) | 70 (68.63) | - |
|  |  | Control | 55 | 1 (1.82) | 54 | 51 (94.44) | 40 (74.07) | - |
| **E12 *in vivo*** | 2 | *NANOG*-targeted | 171 | 22 (12.87) | 149 | 126<br>(78.4 ± 16.7) | 79<br>(43.89 ± 24.74) | - |
|  |  | Control | 142 | 19 (13.38) | 123 | 111<br>(90.7 ± 3.74) | 58<br>(50.08 ± 23.99) | - |
| **E13 *in vivo*** | 1 | *NANOG*-targeted | 99 | 3 (3.03) | 96 | 82 (85.42) | 28 (29.17) | - |
|  |  | Control | 89 | 0 (0.0) | 89 | 80 (89.89) | 34 (38.20) | - |

No statistically significant differences in developmental rates were observed between the *NANOG*-targeted group (microinjected with *NANOG* gRNA and Cas9 mRNA, partially composed of KO embryos) and the control group (microinjected with Cas9 mRNA only, composed exclusively of WT embryos) (mean ± s.e.m.; t-test; $P > 0.05$ for developmental stages with at least three independent replicates; Chi-square test; $P > 0.05$ otherwise). Cleavage and blastocyst rates were calculated based on the number of zygotes. Embryo survival at D12 was calculated from the number of blastocysts transferred to the post-hatching *in vitro* culture system.

### 3.8. Statistical analyses

Statistical analyses were performed using GraphPad Prism (GraphPad Software, San Diego, CA, USA), and significance was set at P ≤ 0.05. Differences in D8 blastocyst rates, D12 embryo survival rates, cell counts at D8 and D12, embryo area and the percentage of hypoblast migration along the inner surface of the embryo at D12, and conceptus length and NANOG+ and SOX2+ cell numbers at E11, E12, and E13 *in vivo* were analysed using a t-test when data followed a normal distribution. When the D’Agostino & Pearson normality test failed, the non-parametric Mann-Whitney test was applied. Differences in epiblast survival, ED formation rates at D12 *in vitro*, and complete hypoblast migration rates and ED formation rates at E11, E12 and E13 *in vivo*, were analysed by Chi-square test.

## 4. Results

### 4.1. Hypoblast cell number is reduced in NANOG KO blastocysts

*NANOG* ablation did not impair embryo development to the blastocyst stage, as cleavage and blastocyst formation rates were comparable between the *NANOG*-targeted group (microinjected with Cas9-encoding mRNA and sgRNA against *NANOG*, thereby partially composed of KO embryos) and the control group (microinjected with Cas9-encoding mRNA aloney and therefore composed exclusively of WT embryos) (**Table 3**). Among the 49 blastocysts analysed from the *NANOG*-targeted group at D8, 33 were classified as KO and lacked NANOG protein expression (**Figure 1**), corresponding to a KO generation efficiency of 67.3%.

**Figure 1.**
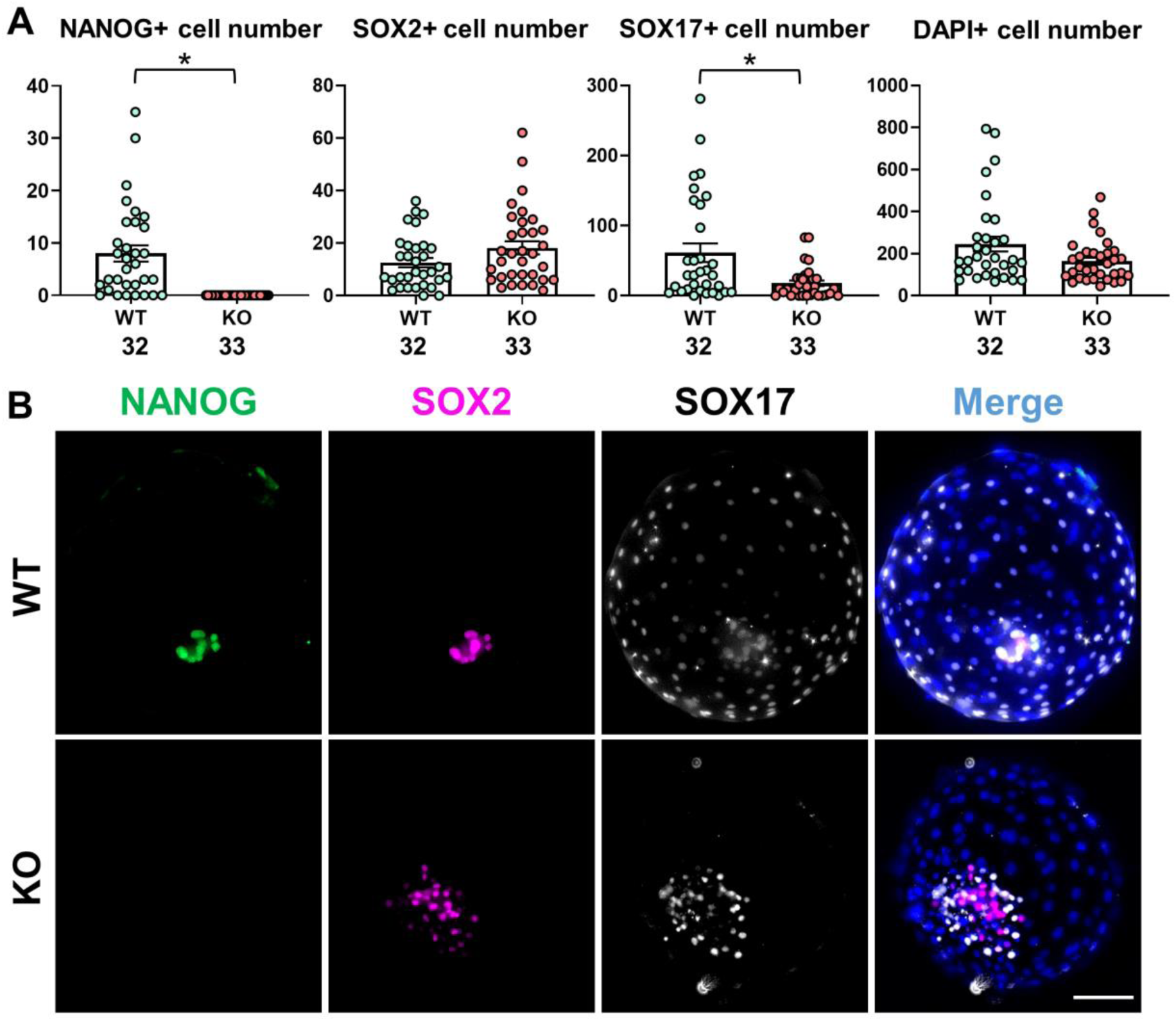
Epiblast and hypoblast differentiation in *NANOG* knockout (KO) and wild-type (WT) day (D) 8 blastocysts. **A)** Scatter plots showing the number of NANOG+, SOX2+, SOX17+ and total cells (mean ± s.e.m.) in *NANOG* KO and WT blastocysts at D8. The number of embryos analysed in each group is shown below each column. Asterisk above columns indicate statistically significant differences (P < 0.05; non-parametric Mann-Whitney test). **B)** Representative immunofluorescence images of *NANOG* KO and WT blastocysts at D8, stained for NANOG (green; epiblast), SOX2 (magenta; epiblast) and SOX17 (white; hypoblast). Nuclei were counterstained with DAPI (merge). Scale bars: 100 µm.

*NANOG* KO blastocysts displayed normal morphology. No significant differences were observed among WT and KO in the number of SOX2+, and SOX17+ cells were detected in most *NANOG* KO blastocysts. However, the number of SOX17+ hypoblast cells was significantly reduced in KO embryos compared with WT embryos (**Figure 1**). These results indicate that *NANOG* ablation does not impair blastocyst formation or early epiblast development, but compromises hypoblast specification or proliferation at the blastocyst stage.

### 4.2. NANOG ablation reduces hypoblast migration during early post-hatching stages in vitro

*NANOG* ablation did not impair early post-hatching development *in vitro*. Embryo survival rates from D6/7 to D12 were comparable between the *NANOG*-targeted and control groups (**Table 3**). Among the 55 embryos analysed from the *NANOG*-targeted group at D12, 37 were KO and lacked NANOG protein expression (**Figure 2**), corresponding to a KO generation efficiency of 67.3%.

**Figure 2.**
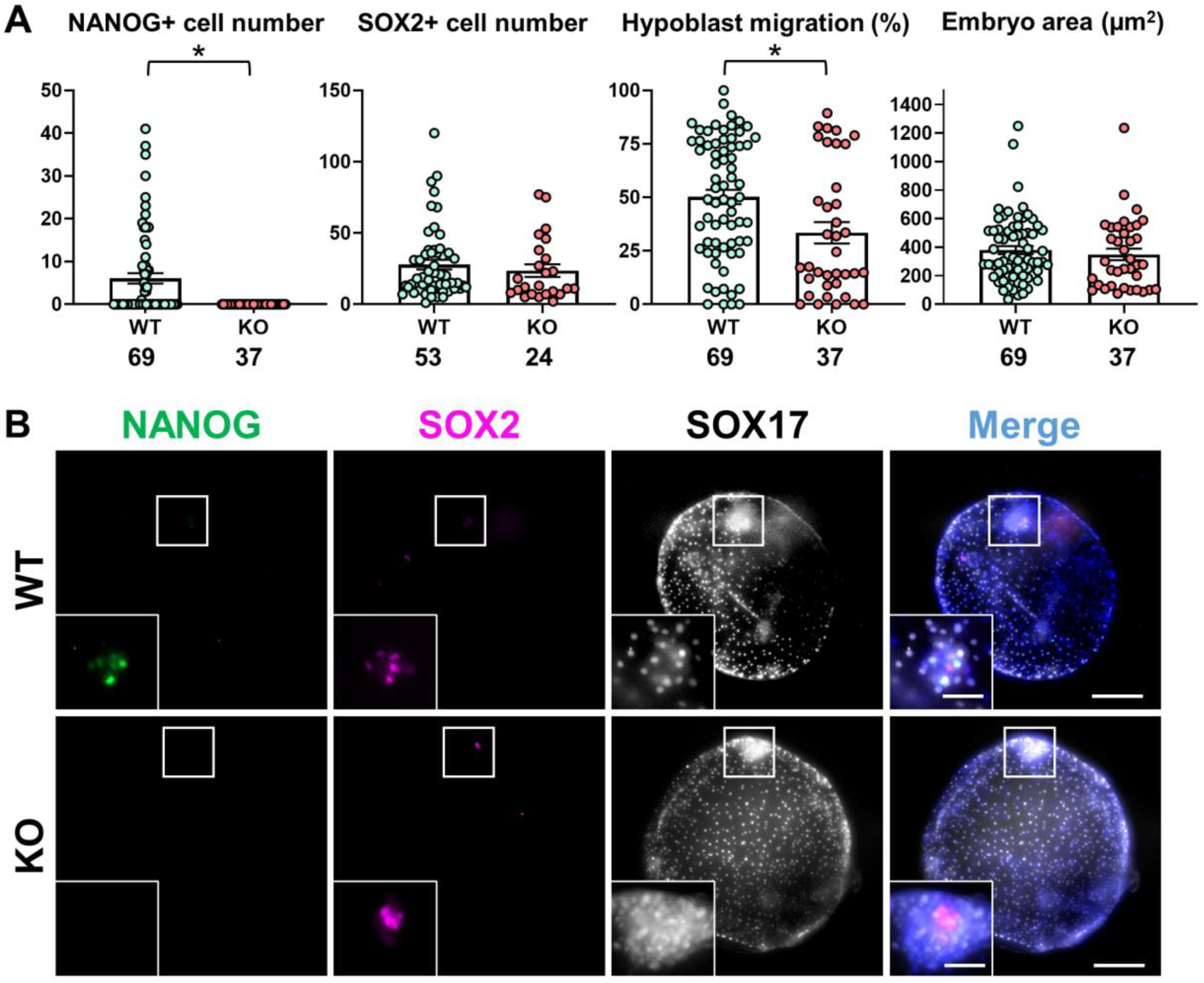
Post-hatching *in vitro* development of *NANOG* knockout (KO) and wild-type (WT) day (D) 12 embryos. **A)** Scatter plots showing the number of NANOG+ and SOX2+ cells, percentage of hypoblast migration and embryo area (mean ± s.e.m.) in *NANOG* KO and WT embryos at D12. The number of embryos analysed in each group is shown below each column. Asterisk above columns indicate statistically significant differences (P < 0.05; non-parametric Mann-Whitney test). **B)** Representative immunofluorescence images of *NANOG* KO and WT embryos at D12, stained for NANOG (green; epiblast), SOX2 (magenta; epiblast) and SOX17 (white; hypoblast). Nuclei were counterstained with DAPI (merge). White boxes in whole embryos indicate magnified views of the embryonic disc (ED) region in the panels below. Scale bars: 200 µm for whole embryos; 50 µm for ED magnifications.

Embryo area did not differ among WT and KO embryos. Similarly, the number of SOX2+ cells (**Figure 2**), the percentage of embryos with surviving epiblast cells (53/69 [76.8%] vs. 24/37 [64.9%] for WT, and KO, respectively), and the percentage of embryos showing ED (19/53 [35.8%] vs. 7/24 [29.2%] for WT and KO, respectively) were not significantly affected by *NANOG* ablation (Chi-square test; P ˃ 0.05).

In contrast, the percentage of hypoblast migration along the inner surface of the TE was significantly reduced in KO embryos compared with WT embryos (**Figure 2**). These findings indicate that *NANOG* ablation reduces hypoblast migration along the inner TE surface during post-hatching development.

### 4.3. NANOG ablation impairs epiblast survival and embryonic disc formation during conceptus elongation

To assess the effects of *NANOG* ablation at later developmental stages, D7 blastocysts from both the *NANOG*-targeted and control groups were transferred to synchronized recipient ewes on embryonic day (E) 6 and conceptuses were recovered seven days later (E13) (**Table 4**). All recovered KO conceptuses lacked NANOG protein expression (**Figure 3A-B**).

**Figure 3.**
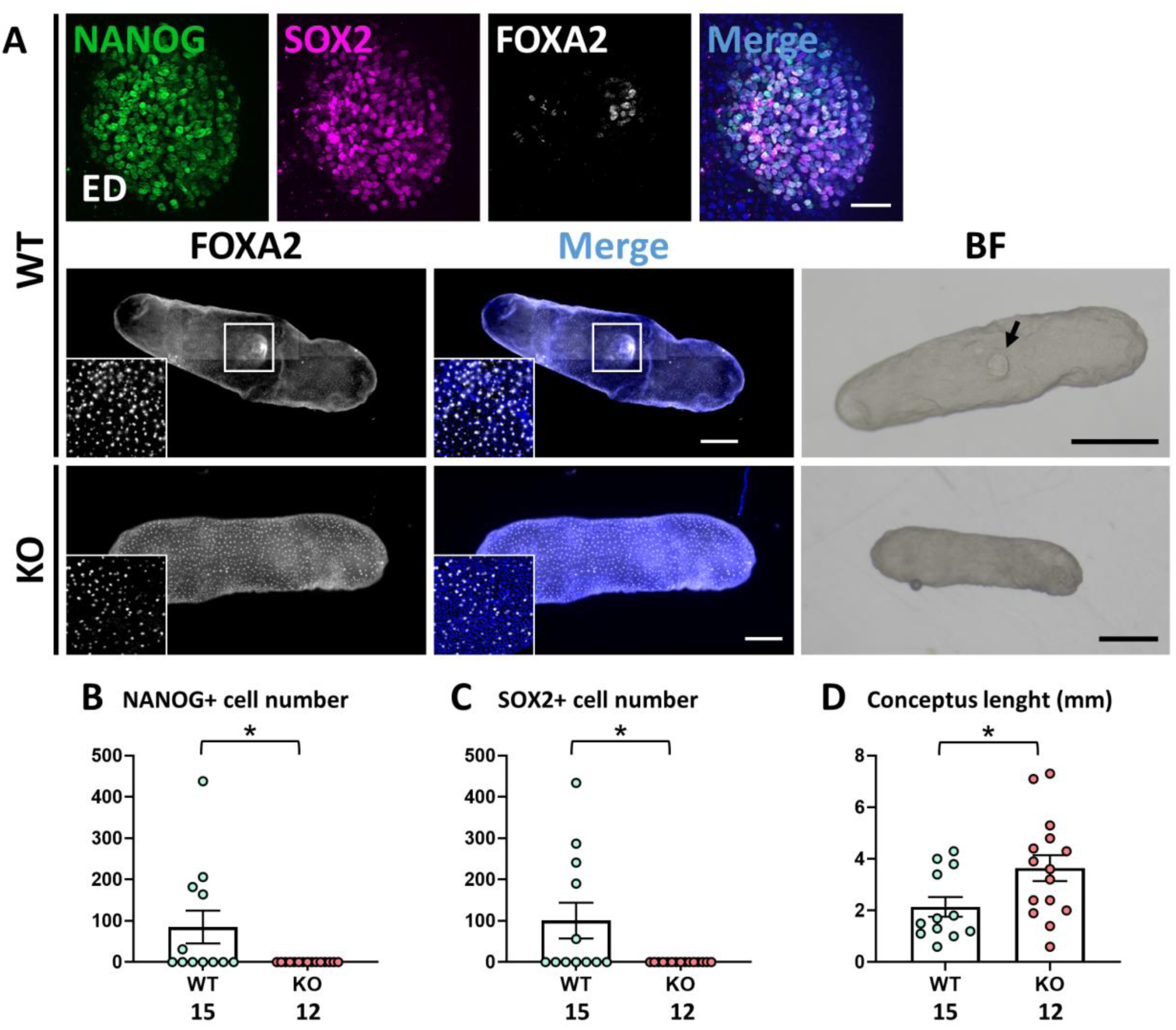
Development of *NANOG* knockout (KO) and wild-type (WT) conceptuses at embryonic day (E) 13. **A)** Representative immunofluorescence and bright field (BF) images of *NANOG* KO and WT conceptuses at E13, stained for NANOG (green; epiblast), SOX2 (magenta; epiblast) and FOXA2 (white; hypoblast). Nuclei were counterstained with DAPI (merge). White boxes in whole embryos indicate magnified views of the embryonic disc (ED) region in the panels above. Insets show magnified regions of hypoblast migration along the inner TE surface. Black arrow indicates the ED. Scale bars: 1 mm for BF pictures; 500 µm for whole embryos; 50 µm for ED magnifications. **B-D)** Scatter plots showing **B)** NANOG+ and **C)** SOX2+ cells and **D)** conceptus length (mean ± s.e.m.) in *NANOG* KO and WT conceptuses at E13. The number of embryos analysed in each group is shown below each column. Asterisk above columns indicate statistically significant differences (P < 0.05; parametric t-test).

**Table 4.**
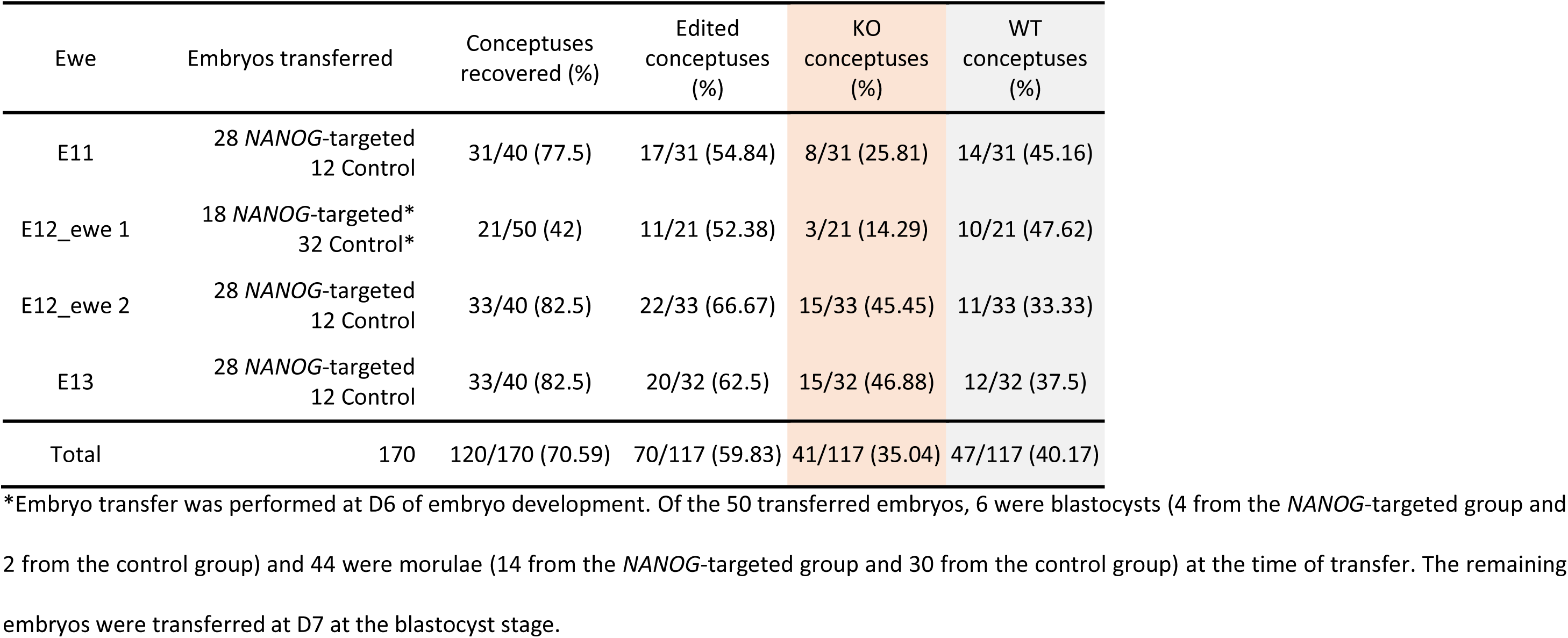
Embryo transfers and CRISPR-Cas9 efficiency. Day (D) 7 blastocysts from the *NANOG*-targeted group (microinjected with *NANOG* gRNA and Cas9 mRNA) and the control group (microinjected with Cas9 mRNA only) were transferred to four recipient ewes. Embryos were recovered at embryonic day (E) 11 (after 5 days), 12 (after 6 days) and 13 (after 7 days) of *in vivo* development. Conceptus recovery rate, editing efficiency, and the proportion of knock-out (KO) and wild-type (WT) embryos are shown.

| Ewe | Embryos transferred | Conceptuses recovered (%) | Edited conceptuses (%) | KO conceptuses (%) | WT conceptuses (%) |
| --- | --- | --- | --- | --- | --- |
| E11 | 28 *NANOG*-targeted<br>12 Control | 31/40 (77.5) | 17/31 (54.84) | 8/31 (25.81) | 14/31 (45.16) |
| E12_ewe 1 | 18 *NANOG*-targeted*<br>32 Control* | 21/50 (42) | 11/21 (52.38) | 3/21 (14.29) | 10/21 (47.62) |
| E12_ewe 2 | 28 *NANOG*-targeted<br>12 Control | 33/40 (82.5) | 22/33 (66.67) | 15/33 (45.45) | 11/33 (33.33) |
| E13 | 28 *NANOG*-targeted<br>12 Control | 33/40 (82.5) | 20/32 (62.5) | 15/32 (46.88) | 12/32 (37.5) |
| Total | 170 | 120/170 (70.59) | 70/117 (59.83) | 41/117 (35.04) | 47/117 (40.17) |

\*Embryo transfer was performed at D6 of embryo development. Of the 50 transferred embryos, 6 were blastocysts (4 from the *NANOG*-targeted group and 2 from the control group) and 44 were morulae (14 from the *NANOG*-targeted group and 30 from the control group) at the time of transfer. The remaining embryos were transferred at D7 at the blastocyst stage.

*NANOG* ablation did not impair the development of EEMs at E13, as no adverse effects were observed in conceptus length and the proportion of conceptuses displaying complete hypoblast migration in KO compared to WT embryos (**Table 5**, **Figure 3A, D**). In contrast, epiblast development was severely compromised in *NANOG*-deficient conceptuses. All KO embryos lacked SOX2+ epiblast cells, whereas 5 of 12 WT embryos (41.7%) displayed compact EDs (**Table 5**, **Figure 3A-C**).

**Table 5.** Hypoblast migration and embryonic disc (ED) development in *NANOG* knock-out (KO) and wild-type (WT) conceptuses.

| Embryonic day | Genotype | Complete hypoblast migration (%) | ED formation (%) |
| --- | --- | --- | --- |
| **E11** | **KO** | 2/8 (25) | 0/8 (0) <sup>a</sup> |
|  | **WT** | 6/14 (42.86) | 8/14 (57.14) <sup>b</sup> |
| **E12** | **KO** | 4/17 (23.53) | 0/18 (0) <sup>a</sup> |
|  | **WT** | 10/21 (47.62) | 13/21 (61.9) <sup>b</sup> |
| **E13** | **KO** | 10/15 (66.67) | 0/15 (0) <sup>a</sup> |
|  | **WT** | 7/12 (58.3) | 5/12 (41.67) <sup>b</sup> |

Different letters indicate statistically significant differences (Chi-square test; $p < 0.05$ ).

To determine the developmental stage at which the ED was lost in *NANOG*-null embryos, conceptuses recovered at E12 and E11 were analysed. The proliferation of EEMs was not significantly affected, as embryo diameter and the proportion of conceptuses displaying complete hypoblast migration were comparable among WT and KO embryos at both E12 and E11 (**Table 5**, **Figure 4D**, **Figure 5D**). In contrast, epiblast development was severely impaired, as no SOX2+ cells were detected in any KO embryo at either E12 or E11 (**Figure 4C**, **Figure 5C**). All KO conceptuses at both developmental stages lacked an ED, whereas ED formation was detected in WT embryos (E12: 13/21 [61.9%]; E11: 8/14 [57.1%]) (**Table 5**, **Figure 4A**, **Figure 5A**).

**Figure 4.**
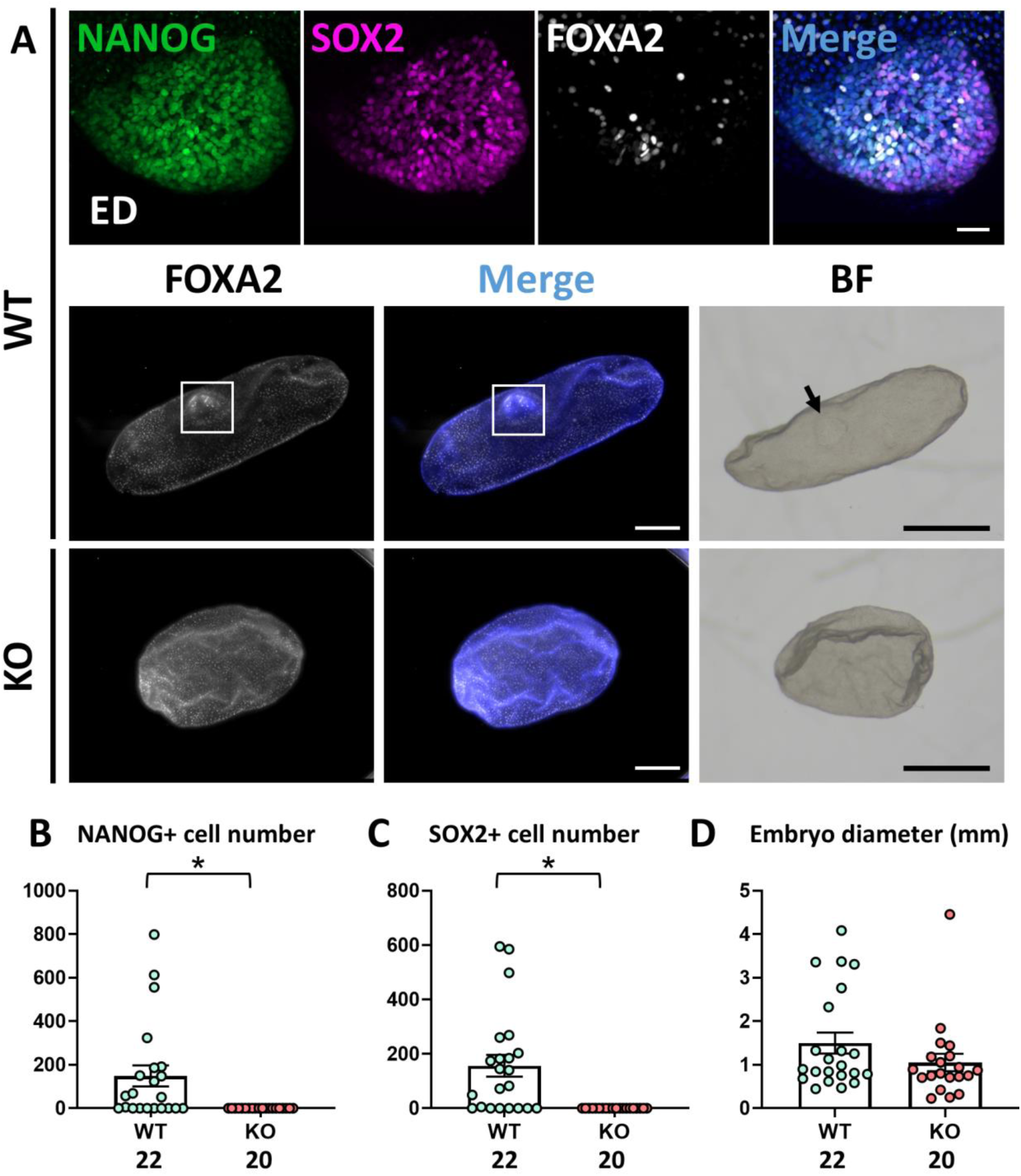
Development of *NANOG* knockout (KO) and wild-type (WT) conceptuses at embryonic day (E) 12. **A)** Representative immunofluorescence and bright field (BF) images of *NANOG* KO and WT conceptuses at E12, stained for NANOG (green; epiblast), SOX2 (magenta; epiblast) and FOXA2 (white; hypoblast). Nuclei were counterstained with DAPI (merge). White boxes in whole embryos indicate magnified views of the embryonic disc (ED) region in the panels above. Black arrows indicate the ED. Scale bars: 1 mm for BF pictures; 500 µm for whole embryos; 50 µm for ED magnifications. **B-D)** Scatter plots showing **B)** NANOG+ and **C)** SOX2+ cells and **D)** embryo diameter (mean ± s.e.m.) in *NANOG* KO and WT conceptuses at E12. The number of embryos analysed in each group is shown below each column. Asterisk above columns indicate statistically significant differences (P < 0.05; non-parametric Mann-Whitney test).

**Figure 5.**
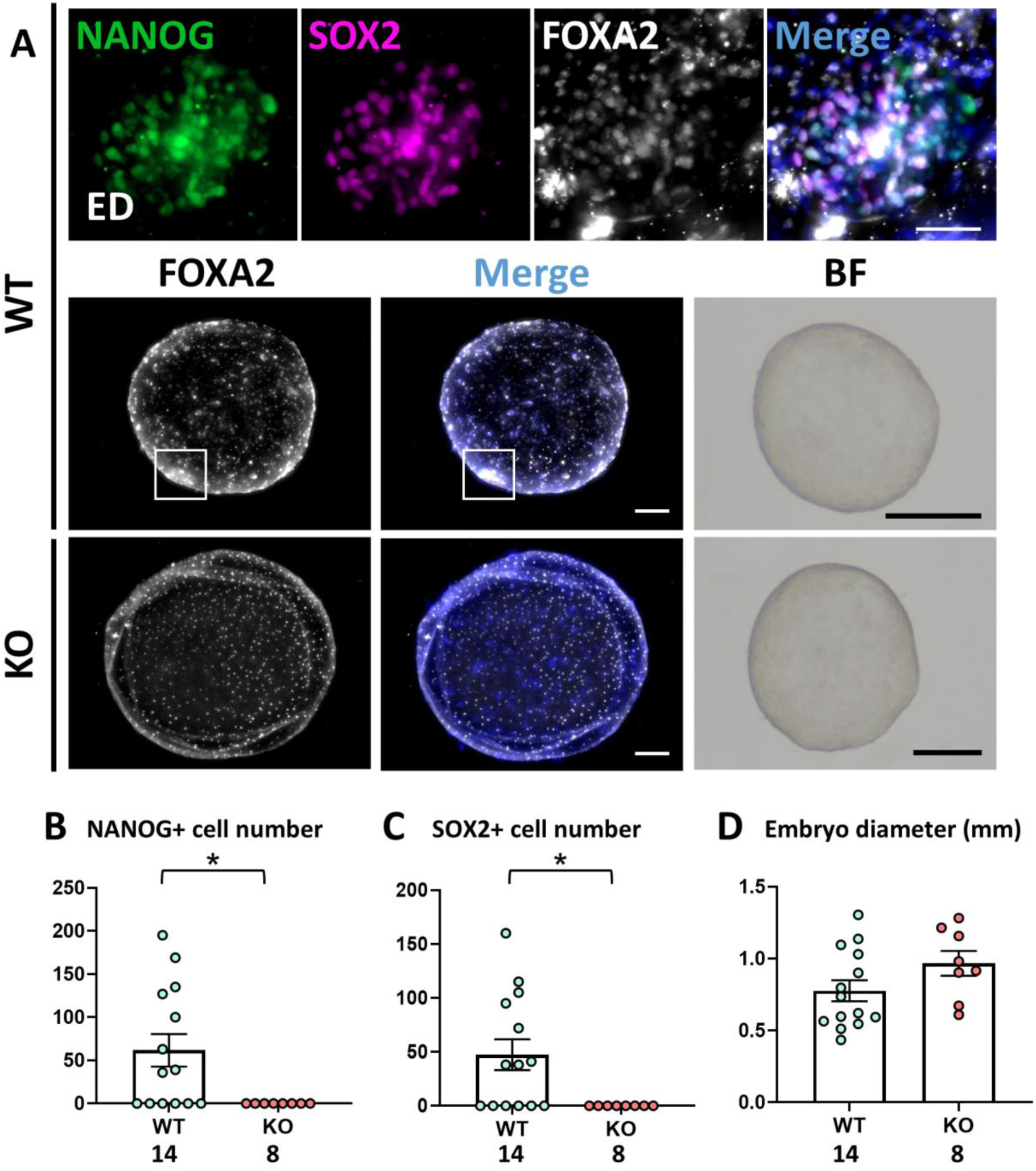
Development of *NANOG* knockout (KO) and wild-type (WT) conceptuses at embryonic day (E) 11. **A)** Representative immunofluorescence and bright field (BF) images of *NANOG* KO and WT conceptuses at E11, stained for NANOG (green; epiblast), SOX2 (magenta; epiblast) and FOXA2 (white; hypoblast). Nuclei were counterstained with DAPI (merge). White boxes in whole embryos indicate magnified views of the embryonic disc (ED) region in the panels above. Scale bars: 500 µm for BF pictures; 200 µm for whole embryos; 50 µm for ED magnifications. **B-D)** Scatter plots showing **B)** NANOG+ and **C)** SOX2+ cells and **D)** embryo diameter (mean ± s.e.m.) in *NANOG* KO and WT conceptuses at E11. The number of embryos analysed in each group is shown below each column. Asterisk above columns indicate statistically significant differences (P < 0.05; non-parametric Mann-Whitney test).

Taken together, these results indicate that early defects in hypoblast specification are compensated during conceptus elongation, whereas epiblast survival is severely compromised in *NANOG* KO embryos, resulting in loss of the ED before E11 *in vivo*.

## 5. Discussion

In this study, we explored the role of *NANOG*, an epiblast-specific transcription factor conserved across mammals, during early sheep embryo development. Our results demonstrate that *NANOG* plays stage-dependent roles during ovine embryogenesis. Although *NANOG* ablation did not compromise blastocyst formation or the establishment of the early epiblast, it reduced the number of hypoblast cells at the blastocyst stage and hypoblast migration during early post-hatching development. These detrimental effects on the hypoblast were not detected at later stages, as *NANOG* KO conceptuses recovered *in vivo* displayed hypoblast development comparable to WT conceptuses, thereby indicating a compensatory mechanism. In contrast, *NANOG* deficiency resulted in the complete loss of epiblast cells E11.

In line with previous studies in mouse (Bower *et al*., 2026, Frankenberg *et al*., 2011, Messerschmidt and Kemler, 2010, Mitsui *et al*., 2003), human (Bower *et al*., 2026), porcine (Lee *et al*., 2023), and bovine (Ortega *et al*., 2020, Springer *et al*., 2021) embryos, *NANOG* was dispensable for blastocyst formation in sheep. At this stage, an essential role for NANOG on hypoblast specification has been reported in the mouse, where, although ICM cells in *Nanog* KO embryos express the early hypoblast marker GATA6, expression of later hypoblast markers, including SOX17 and GATA4, is totally impaired (Bower *et al*., 2026, Frankenberg *et al*., 2011, Messerschmidt and Kemler, 2010). The effect of *NANOG* ablation on hypoblast development in bovine and porcine embryos remains less clear, with conflicting data regarding the effect on GATA6 positive cells (Lee *et al*., 2023, Ortega *et al*., 2020, Springer *et al*., 2021). However, GATA6 is not a specific hypoblast marker in ungulate species at the blastocyst stage (Martínez de Los Reyes *et al*., 2024, Perez-Gomez *et al*., 2021). We therefore used SOX17 as a more specific marker of hypoblast identity and found that SOX17 expression was retained in most *NANOG* KO blastocysts. This observation is consistent with previous studies reporting SOX17 expression in bovine (Springer *et al*., 2021), porcine (Lee *et al*., 2023), and human (Bower *et al*., 2026) *NANOG* KO blastocysts, suggesting that, unlike in the mouse, NANOG is not required for hypoblast specification in non-rodent mammals.

Despite the dispensable role of NANOG on hypoblast specification, our results indicate that NANOG contributes to some extent to early hypoblast development in sheep. *NANOG* KO blastocysts contained significantly fewer SOX17+ cells than WT embryos, and this was followed by reduced hypoblast migration along the inner surface of the TE during post-hatching development up to D12 *in vitro*. We recently observed a similar phenotype in bovine *NANOG* KO embryos following extended *in vitro* culture, in which the proportion of embryos displaying complete hypoblast migration was reduced at D12 (Flores-Borobia *et al*., 2026b). However, when *NANOG* KO sheep conceptuses were recovered following *in vivo* development at E11-13, hypoblast development was comparable to that of WT conceptuses. A similar observation was made in bovine embryos, in which all *NANOG* KO and WT conceptuses recovered at E14 displayed complete hypoblast migration (Flores-Borobia *et al*., 2026b). These findings suggest that the early defects in hypoblast development associated to *NANOG* ablation can be compensated during subsequent conceptus elongation in ungulates. The apparent recovery of hypoblast development *in vivo* may also highlight the contribution of the uterine environment, which could provide signals or cellular interactions that are absent or insufficient under *in vitro* culture conditions (Siegmund-Sabater *et al*., 2026).

The role of NANOG in pluripotency and epiblast development has been extensively investigated in the mouse (Chambers *et al*., 2007, Messerschmidt and Kemler, 2010, Mitsui *et al*., 2003, Sun *et al*., 2014). In *Nanog* KO embryos, NANOG loss had no apparent effect on the number of OCT4+ cells in E3.5 blastocysts, whereas a reduction in OCT4+ cells was observed at later stages by E4.5 (Messerschmidt and Kemler, 2010). Similarly, the comparable number of SOX2+ cells in sheep *NANOG* KO and WT D8 blastocysts suggest that NANOG is not required for maintaining the initial pluripotency of the inner cell mass in sheep. This is consistent with a study in bovine embryos reporting similar numbers of SOX2+ cells in *NANOG* KO and WT D8 blastocysts (Springer *et al*., 2021), and with another study in human embryos in which *SOX2* expression levels were not significantly reduced in *NANOG*-edited compared with WT D6 blastocysts (Bower *et al*., 2026). However, other studies have reported reduced *SOX2* mRNA levels in D8 bovine blastocysts (Ortega *et al*., 2020) and reduced SOX2+ cells in D7 porcine blastocysts (Lee *et al*., 2023). These discrepancies may reflect differences in experimental settings and genotyping strategies.

Contrasting with the dispensable role of *NANOG* in early pluripotency, all *NANOG* KO conceptuses lacked SOX2+ epiblast cells and an ED. Similarly, no EDs were detected in *NANOG* KO bovine conceptuses recovered at E14 (Flores-Borobia *et al*., 2026b). Thus, the requirement for NANOG in pluripotency maintenance in ungulates seems to become particularly important around the onset of ED formation, when the initially unorganized population of epiblast cells undergoes polarization to form a columnar epithelium (Martínez de los Reyes *et al*., 2026). A similar temporal pattern has been described in the mouse, where the consequences of *Nanog* loss become apparent after the blastocyst stage, from E4.5 (Messerschmidt and Kemler, 2010, Mitsui *et al*., 2003) to E5.5 (Sun *et al*., 2014), coinciding with the transition of the mouse epiblast from an unpolarised cluster of pluripotent cells into a hollow, cup-shaped columnar epithelium (Bedzhov *et al*., 2014, Kim *et al*., 2021). Although the morphology and timing of epiblast morphogenesis differ between rodents and ungulates, these observations raise the possibility of a conserved requirement for NANOG during the transition from an early pluripotent epiblast or inner cell mass to an organized epithelial embryonic structure. Further functional studies would be required to confirm this hypothesis in other species, in which current studies are limited to the blastocyst stage.

The ability of the extraembryonic membranes, including the TE and hypoblast, to proliferate and continue development in the absence of an ED in *NANOG* KO embryos further supports previous observations that conceptus elongation is uncoupled from ED development (Flores-Borobia *et al*., 2024, Flores-Borobia *et al*., 2026b). This finding has important implications for early pregnancy assessment in livestock, as current early pregnancy detection methods rely on signals derived exclusively from the TE (Filho *et al*., 2020, Kizaki *et al*., 2013, Roberts *et al*., 2015) and therefore cannot distinguish between real pregnancies with conceptuses carrying an ED and anembryonic pregnancies (Flores-Borobia *et al*., 2026a). Consequently, the identification of the molecular mechanisms that specifically regulate ED development, independently of extraembryonic membranes growth, may contribute to a better understanding of early pregnancy loss. This is particularly relevant given the high incidence of early embryonic loss in both humans (Larsen *et al*., 2013, Macklon *et al*., 2002) and livestock species (Almeida and Dias, 2022, Diskin and Morris, 2008, Ealy *et al*., 2019, Flores-Borobia *et al*., 2026a, Wilmut *et al*., 1986, Wiltbank *et al*., 2016). Mechanistic studies in non-rodent species, now increasingly feasible through genome-editing approaches, may therefore provide important insights into the mechanisms underlying these early developmental failures.

Overall, our findings identify NANOG as a key regulator of post-hatching epiblast maintenance and ED development in sheep. While NANOG is not required for initial epiblast pluripotency, it becomes essential for the maintenance and developmental progression of the epiblast during ED formation. This requirement appears to be conserved across rodents, ungulates and primates. In contrast, NANOG loss causes an early reduction in hypoblast cell numbers and impairs hypoblast migration *in vitro*, but these defects are compensated at later stages. These findings indicate that NANOG is not essential for hypoblast development in sheep, consistent with previous studies in bovine and human embryos and in contrast to the mouse. This further supports the notion that a significant number of the mechanisms regulating early lineages development are mouse-specific and highlights the importance of studying these processes directly in non-rodent mammals. Our results therefore expand our understanding of the molecular mechanisms governing lineage development during post-hatching stages in non-rodent mammals.

## 6. Acknowledgements

The authors want to acknowledge the slaughterhouse Matadero Mondejano S. L. and the veterinarians for kindly providing ovine ovaries for the experiments.

## 7. Declaration of interest

There is no conflict of interest that could be perceived as prejudicing the impartiality of the research reported.

## 8. Grant Support

This work has been funded by the projects PID2021-122153NA-I00 and PID2024-155682NB-I00 from the Spanish Ministry of Science and Innovation to PRI, and ECQ2018-005184-P from the Spanish Ministry of Science and Innovation to PBA. MCS was funded by a Margarita Salas fellowship.

## 9. Author contributions

NMR, MCS, PM and PRI contributed to ovine *in vitro* embryo production and IF analyses. PBA designed and produced CRISPR-Cas9 components. Embryo genotyping was carried out by NMR. ATD, JSM and PBA were involved in embryo transfer experiments. PRI and PBA provided protocols, resources and supervised the team. ATD, JSM and PBA performed embryo transfers. NMR and PRI drafted the manuscript, and all authors reviewed it. PRI and PBA acquired funding.

## 10. Data availability statement

The data underlying this article are available in the article.

## Notes

### Competing Interest Statement

The authors have declared no competing interest.

